# PocketMaster: a flexible, automated tool for analyzing, clustering, and visualizing protein pocket structural diversity

**DOI:** 10.1101/2025.10.30.685483

**Authors:** Narek Abelyan

**Author notes:** Laboratory of Computational Modeling of Biological Processes, Institute of Molecular Biology, National Academy of Sciences of the Republic of Armenia (NAS RA), 0014, Yerevan.

## Abstract

PocketMaster is a flexible and automated tool for the analysis, clustering, and interpretation of protein pockets. It enables the exploration of structural diversity in functional and interacting regions of proteins, supports scalable alignment, RMSD calculation, and hierarchical clustering of hundreds to thousands of models. The tool provides multiple pocket-detection methods, alignment algorithms, and clustering approaches, allowing analyses to be tailored to specific scientific questions. With these capabilities, PocketMaster can be particularly valuable in the early stages of drug design, where accurate analysis and selection of protein structures are essential. Using TYK2 as a case study, PocketMaster demonstrated its ability to identify conformational differences between kinase and pseudokinase domains, as well as subtypes of structures within each domain, reflecting the influence of various ligands and protein states. The results confirm known structural features and illustrate the potential of the tool for systematic exploration of protein pockets, quantitative assessment of differences, and support of rational drug design. The PocketMaster source code, together with example input files and documentation, can be accessed at https://github.com/narek-abelyan/PocketMaster

## INTRODUCTION

The three-dimensional structure of a protein is not a static entity, but a dynamic system capable of adopting multiple conformational states (1). Particularly important are the regions involved in the specific binding of small molecules or proteins - active sites, allosteric pockets, catalytic regions, and PPI pockets (2). Even minor changes in these areas can significantly affect protein function. Conformational variations can be induced by various factors, including ligand binding, allosteric effects, changes in pH, temperature, or point mutations, especially if they occur near a functional site (3, 4).

Analyzing such changes is a key step in understanding the molecular mechanisms of protein function, as well as in selecting the optimal structure and conformation for drug development targeting specific protein states (5). Comparing the pockets of the same protein under different conditions allows a deeper exploration of its structural diversity, revealing patterns that stabilize interactions or, conversely, uncovering mechanisms that disrupt function - for example, in the case of disease-related mutations. This can be particularly useful in the early stages of drug development, when precise analysis and selection of suitable protein structures are critical. Thousands of structures reflecting different functional states, mutant forms, and interactions with various ligands or inhibitors are already available in the PDB (6). Nevertheless, there is still no universal, convenient, and automated tool for comprehensive analysis of such structural changes at the level of local functionally relevant regions of a protein, with the capacity for quantitative assessment and clear visualization of differences.

To address this challenge, we developed PocketMaster - a flexible and scalable software solution for analyzing structural similarities between protein pockets. Implemented in Python and integrated with the PyMOL API (7), PocketMaster provides automated processing of large structural datasets, including alignment, pocket construction, RMSD calculation, and visualization of results. The tool is designed to efficiently handle hundreds to thousands of protein models, enabling comprehensive, quantitatively supported, and visually interpretable comparative analyses.

PocketMaster provides a broad range of flexible settings, enabling analyses to be tailored to diverse research objectives and structural data types. The tool supports multiple approaches for pocket identification, a variety of RMSD calculation methods, and clustering algorithms, allowing researchers to select optimal parameters for each specific study. Additionally, PocketMaster automatically saves results and generates informative visualizations and reports. This comprehensive functionality ensures convenience, scalability, and highly interpretable analysis of protein pockets, facilitating an in-depth examination of their structural features and patterns across large datasets.

## MATERIALS AND METHODS

### Main Functionalities of PocketMaster

PocketMaster offers a high degree of flexibility and configurability, allowing it to be adapted for a wide range of research tasks. The main customization features include:

#### 1. Importing Protein Structures

PocketMaster supports several modes for obtaining three-dimensional protein structures:

- Local PDB files: Users can load files from a specified folder, which is convenient when working with proprietary or pre-processed data.
- UniProt ID: The software can automatically retrieve and load all available PDB structures corresponding to a given UniProt ID (8).
- PDB ID: PocketMaster can identify the corresponding UniProt ID from a PDB ID and load all associated PDB structures.

#### 2. Flexible Preprocessing of Structures

Before analysis, PocketMaster offers preprocessing, which can help standardize the data and ensure accurate structural comparisons. This includes removal of water molecules, ions (e.g., Mg^2+^, Cl^−^), sulfates, phosphates (SO_4_, PO_4_), buffer components (TRS, MES, HEPES), cryoprotectants (GOL, EDO, MPD), reducing agents (DTT, BME, TCEP), modified amino acid residues (CSO, MSE, SEP, TPO, PTR), alternate conformations (ALTLOC), hydrogen atoms, anisotropic parameters (ANISOU), and other elements not belonging to the main polymer chain.

#### 3. Multiple Approaches for Defining Alignment Regions

PocketMaster provides multiple methods for defining the alignment region and subsequent RMSD calculation, ensuring precise selection of functionally relevant areas. This feature is particularly important for analyzing complex or non-standard protein structures (Fig. 1).

**Figure 1:**
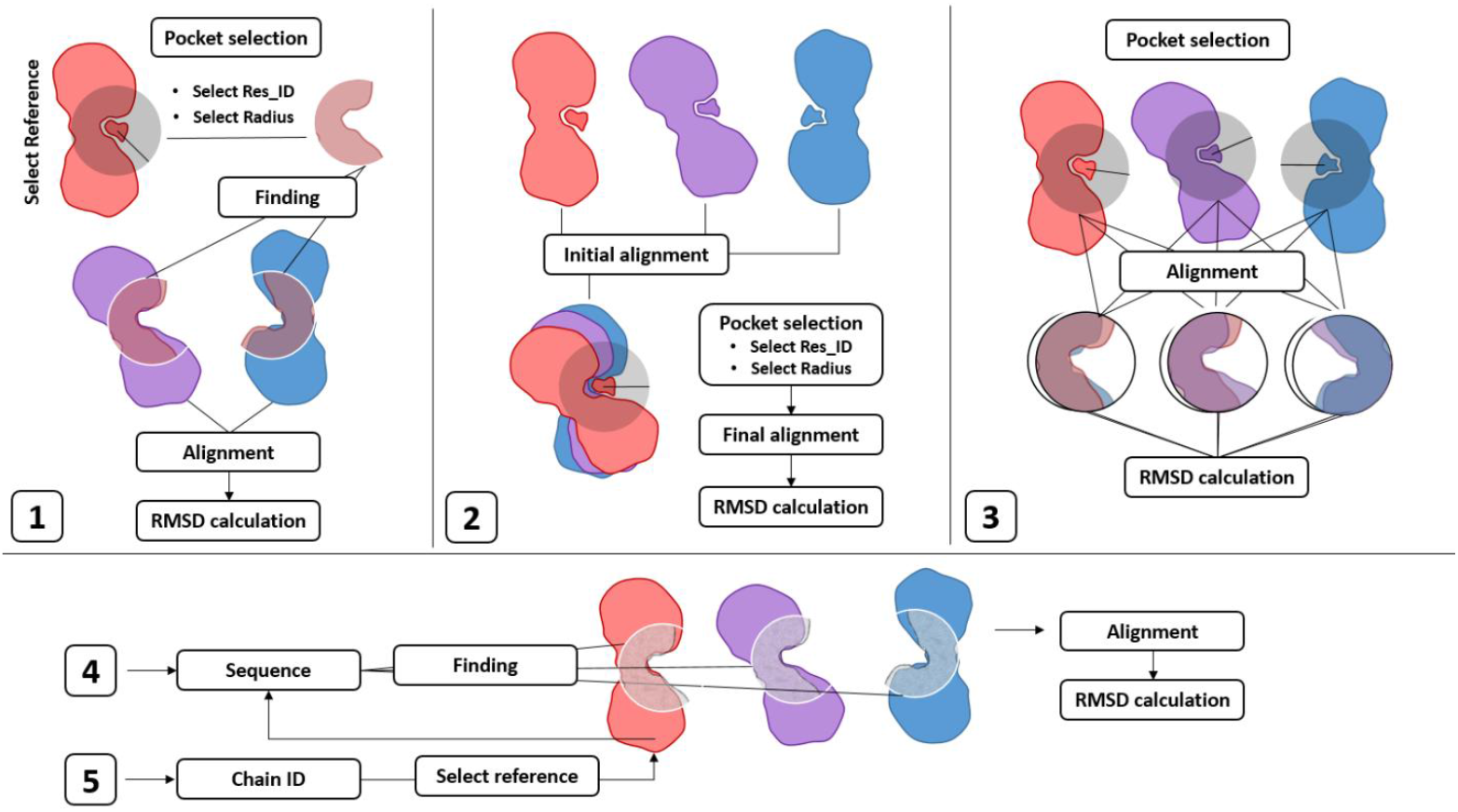
Supported methods for defining alignment regions and their detailed interpretation.

1. The region is defined on the selected reference structure by a specified residue ID and radius (Å), and then this region is searched for and aligned in all structures.
2. The region is defined after a preliminary alignment of all structures to the reference, for each structure around the selected residue within a given radius (Å).
3. The region is defined for each structure around its HET groups within a specified radius (Å). With proper preprocessing, it is possible to keep only ligands in the HET groups.
4. The region is defined according to a user-provided list of residues, and then this region is searched for and aligned in all structures.
5. The region is defined on the reference structure by a specified chain identifier, and then this region is searched for and aligned in all structures.

#### 4. Alignment and RMSD calculation methods

PocketMaster supports multiple algorithms for structural alignment and RMSD calculation, each suitable for different analysis scenarios. Users can select the most appropriate method depending on the task, required accuracy, the presence of unmatched atoms between structures, and computational efficiency:

1. align – A strict alignment method that matches residues according to the amino acid sequence and performs an optimal superposition of the moving structure onto the target. Atoms without a corresponding match between structures are excluded from the calculation. RMSD is then computed based on the matched atoms. Atoms without correspondence or those deviating strongly (beyond the default cutoff of 2 Å) may be excluded from the RMSD calculation. This method is suitable for high-accuracy similarity assessments of structures with similar sequences.
2. cealign – A structural alignment method based on the Combinatorial Extension (CE) algorithm. It aligns protein structures by the geometry of their fragments, minimizing the RMSD between matching Cα atoms. In the final stage of the global alignment, RMSD is calculated only for the Cα atoms of matched segments, while other atoms and non-matching residues are ignored in the calculations, although they move together with the moving structure. This method is particularly effective for comparing structures with different sequences, varying conformations, or mismatched individual atoms.
3. super – A more flexible algorithm, a modified alignment method similar to align, but more robust to sequence mismatches and residue numbering differences. It performs structural matching while accounting for insertions, deletions, and partial mismatches, providing more reliable alignments when there are moderate differences between structures.
4. rms – A direct RMSD calculation between two structures without alignment, assuming a complete one-to-one correspondence of atoms in order. This method is sensitive to mismatches and does not allow differences in structural composition. It is used to assess identical or strictly fitted structures when preliminary alignment has already been performed.
5. rms_cur – A simplified and faster method similar to rms, but with lower computational overhead. It assumes complete atom correspondence and a fixed sequence. This method is suitable for screening large numbers of structures when a quick, though less precise, assessment is required.

Each method is implemented using PyMOL’s built-in tools, ensuring reproducibility of results and compatibility with standard practices in structural analysis.

#### 5. RMSD calculation over all atoms or only Cα atoms

Users can choose to compute RMSD using either all atoms or only α-carbon atoms, allowing flexibility in the accuracy and sensitivity of similarity assessments depending on the specific task and the quality of the data.

#### 6. Hierarchical clustering methods

PocketMaster implements several agglomerative hierarchical clustering algorithms, allowing protein sites to be grouped based on their structural similarity (expressed as an RMSD distance matrix). Users can select the most suitable method depending on the nature of the data and the research objectives. All algorithms are implemented using the SciPy library (9) and rely on a standard distance metric (typically RMSD calculated after alignment):

1. ward – Merges clusters in a way that minimizes differences within each group. It works best with Euclidean distances (e.g., RMSD) and produces compact clusters that are balanced in size. This method is particularly useful when it is important for structures within the same cluster to be as similar as possible.
2. Single – Uses the minimum distance between elements of different clusters (also known as the nearest-neighbor method). It can lead to elongated clusters with a linear structure and is highly sensitive to outliers. Particularly useful for sequentially merging similar structures when tracking gradual changes is important.
3. complete – Merges clusters based on the largest distance between their elements (farthest-neighbor method). This results in more compact and clearly separated groups. This method is suitable when it is important to reliably separate different clusters. It is also less sensitive to outliers but can sometimes merge fairly dissimilar structures prematurely.
4. average (UPGMA) – Calculates the distance between clusters as the average of all pairwise distances between elements from the two clusters. Positioned between the single method (minimum distance) and the complete method (maximum distance), it provides a more balanced and even division of groups. This approach is often used for constructing evolutionary and structural trees.
5. centroid – Measures the distance between clusters based on their centroids, i.e., the mean positions of all points in each cluster. This approach usually gives clear results but can sometimes produce incorrect clustering if non-standard distance measures are used, potentially resulting in an unordered cluster tree.
6. median – Similar to centroid but uses the median instead of the mean. More robust to outliers, though less commonly used in bioinformatics.
7. weighted (WPGMA) – A simplified variant of the average method, where the “weight” (importance) of clusters does not change during merging. This method is used less frequently but can be useful when the data has a known hierarchical structure and it is important to preserve the influence of the original groups without distortion during merging.

All methods are compatible with customizable clustering stop criteria (by number of clusters, distance threshold, or automatic estimation), making the analysis highly flexible.

#### 7. Clustering parameters

PocketMaster provides three main options for forming clusters:

1. By number of clusters (maxclust) - The user specifies the desired number of clusters in advance. The algorithm merges elements until the specified number is reached. This approach is useful when the approximate structure of the data is known or when a specific number of groups is required for further analysis.
2. By distance threshold (distance) - Clusters are formed until the distances between groups exceed a specified threshold. This allows control over intra-cluster similarity: the lower the threshold, the more homogeneous the groups. This method is particularly suitable when clearly defined boundaries between clusters are important.
3. Automatic mode - The algorithm determines the optimal number of clusters based on the data and predefined quality criteria. This mode is useful when the number of clusters is unknown or when the goal is to obtain the most informative grouping without manual tuning. Two approaches are available for automatic clustering:
  a. Elbow method - The number of clusters is selected at the “elbow” point of a plot showing within-cluster distance versus number of merges, where additional clusters result in minimal improvement in compactness.
  b. Threshold based on 70 % of the maximum merge distance - Clusters are formed until the distance between merging groups exceeds 70 % of the maximum observed distance, allowing identification of naturally homogeneous groups.

The ability to choose the clustering method provides greater flexibility and allows the analysis to be adapted to specific tasks, improving the interpretation of results and the quality of the identified groups.

#### 8. Saving Preprocessed, Aligned Structures and Analysis Results

Saving allows easy storage, viewing, and reuse of data, simplifying the integration of PocketMaster into research workflows and ensuring reproducibility.

#### 9. Generating concise reports on structural and sequence differences of pockets

The reports provide key information on variations in the amino acid composition of selected structural regions, facilitating their comparison and interpretation.

#### 10. Creating Clear Visualizations: Heatmaps, Dendrograms, and Histograms

Visualizations help identify recurring patterns, group similar structures, and quickly assess overall similarities and differences.

In summary, PocketMaster offers researchers a wide range of configurable options, enabling accurate and comprehensive analysis of protein sites, while allowing the workflow to be adapted to specific scientific tasks and data volumes.

### Program launch modes and settings

PocketMaster offers two main modes of operation. In the interactive mode, users enter analysis parameters manually via the console, allowing flexible real-time adjustment of the task. To run the program in this mode, the following command is used:

python PocketMaster.py

The second mode uses a YAML configuration file, where all necessary parameters are predefined. This ensures automation and reproducibility of the analysis, especially when working with large datasets or when repeated runs with identical settings are required. The program is launched with a configuration file using the command:

python PocketMaster.py --config config.yaml

When running in interactive mode, a run_config.txt file is generated at the end. It automatically saves all parameters entered by the user, allowing you to run the script later in automatic mode with the same settings, without re-entering parameters. This approach enhances the convenience and efficiency of using PocketMaster in various research scenarios.

## RESULTS

### Application of PocketMaster on TYK2: Analysis of Ligand-Binding Pocket Variability

To demonstrate the capabilities and practical application of PocketMaster, we performed a structural analysis of the TYK2 protein (Tyrosine kinase 2) - a member of the Janus kinase family that plays a crucial role in cytokine signaling and immune regulation (10, 11, 12). TYK2 is considered a promising therapeutic target for autoimmune disorders such as psoriasis, rheumatoid arthritis, and systemic lupus erythematosus (10, 13, 14). In recent years, efforts to develop selective TYK2 inhibitors have resulted in a large number of crystal structures of the protein in complex with various ligands (12, 15, 16, 17), making TYK2 a good subject for analyzing the conformational diversity of its ligand-binding pocket.

The starting dataset was built using UniProt ID: P29597, from which all available TYK2 crystal structures were automatically retrieved from the Protein Data Bank (PDB). In total, 50 structures were collected, differing in bound ligand type, activation state, and, in some cases, the presence of point mutations.

To standardize the input data, preprocessing was performed (do_preprocess=1), which included the removal of water molecules (clean_options:1 - water), ions (clean_options:2 - MG, CL, etc.), sulfates and phosphates (clean_options:3 - SO4, PO4, etc.), buffer components (clean_options:4 - TRS, MES, etc.), cryoprotectants (clean_options:5 - GOL, EDO, etc.), reducing agents (clean_options:6 - DTT, BME, TCEP), alternative atom conformations (clean_options:10 - altloc), and anisotropic parameters (clean_options:11 - anisou). This step removed structural artifacts and improved the quality of subsequent alignments.

All structures were preliminarily aligned to a reference structure (ref_align = 6NZP, init_align = 1, init_align_chain_id = “A”). The reference structure chosen for alignment was PDB ID: 6NZP, containing the inhibitor LB7 tightly bound in the active site. The binding pocket was defined individually for each structure (pocket_method:2) around the reference ligand LB7 (ligand_resi: 901, ligand_chain: A) using a 7 Å radius (radius:7), covering the entire ligand-binding region and including all residues potentially involved in ligand recognition and stabilization.

To evaluate the structural similarity of binding pockets across all TYK2 structures, an all-vs-all comparison was performed (comparison_mode:1). The RMSD values were calculated using the super algorithm (rmsd_method:2 - super), which allows flexible alignment even in the presence of partial atom mismatches between compared structures. The procedure generated a complete RMSD distance matrix representing the pairwise pocket dissimilarities among all structures.

Subsequently, hierarchical clustering was applied using the average linkage (UPGMA) method (linkage_method:4 - average (UPGMA)), while the optimal clustering threshold was automatically determined via the elbow method (cl_choice:3 - Elbow method).

This workflow enabled the identification of distinct groups of models sharing high conformational similarity.

Clustering results were visualized as distance distribution histograms, an RMSD heatmap, and a dendrogram. According to the results, all TYK2 structures were clearly classified into two main classes, which differ significantly in conformation (Figure 2, blue and red clusters on the heatmap). These pronounced structural differences are fully consistent with known literature and scientific data, reflecting the presence of two TYK2 domains – kinase and pseudokinase – which differ substantially in their spatial organization (12, 16, 18).

**Figure 2:**
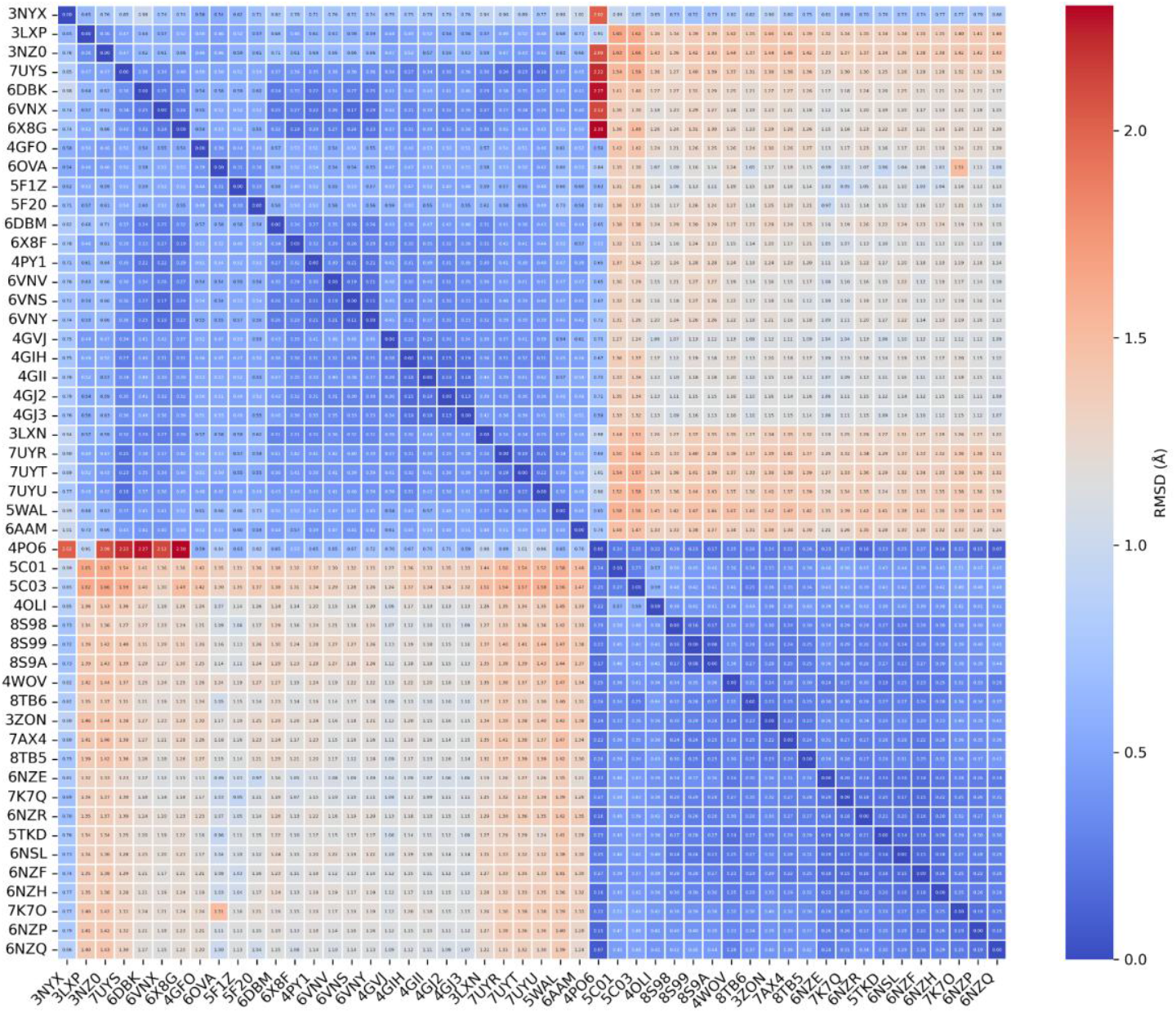
Ordered heatmap representing the full RMSD matrix across all TYK2 structures.

Structures with PDB IDs: 3NYX, 3LXP, 3NZ0, 7UYS, 6DBK, 6VNX, 6X8G, 4GFO, 6OVA, 5F1Z, 5F20, 6DBM, 6X8F, 4PY1, 6VNV, 6VNS, 6VNY, 4GVJ, 4GIH, 4GII, 4GJ2, 4GJ3, 3LXN, 7UYR, 7UYT, 7UYU, 5WAL, 6AAM, and 4PO6 represent kinase domains.

Structures with PDB IDs: 5C01, 5C03, 4OLI, 8S98, 8S99, 8S9A, 4WOV, 8TB6, 3ZON, 7AX4, 8TB5, 6NZE, 7K7Q, 6NZR, 5TKD, 6NSL, 6NZF, 6NZH, 7K7O, 6NZP, and 6NZQ contain the pseudokinase domain.

It is the latter, pseudokinase domain, that binds deucravacitinib, a recently developed first-in-class allosteric TYK2 inhibitor (19, 20). Additionally, the data presented in the RMSD histograms and the hierarchical clustering dendrogram further support this conclusion (Figures 3 and 4). They clearly show the separation of structures into two distinct clusters, highlighting the fundamental differences between the kinase and pseudokinase domains of TYK2.

**Figure 3:**
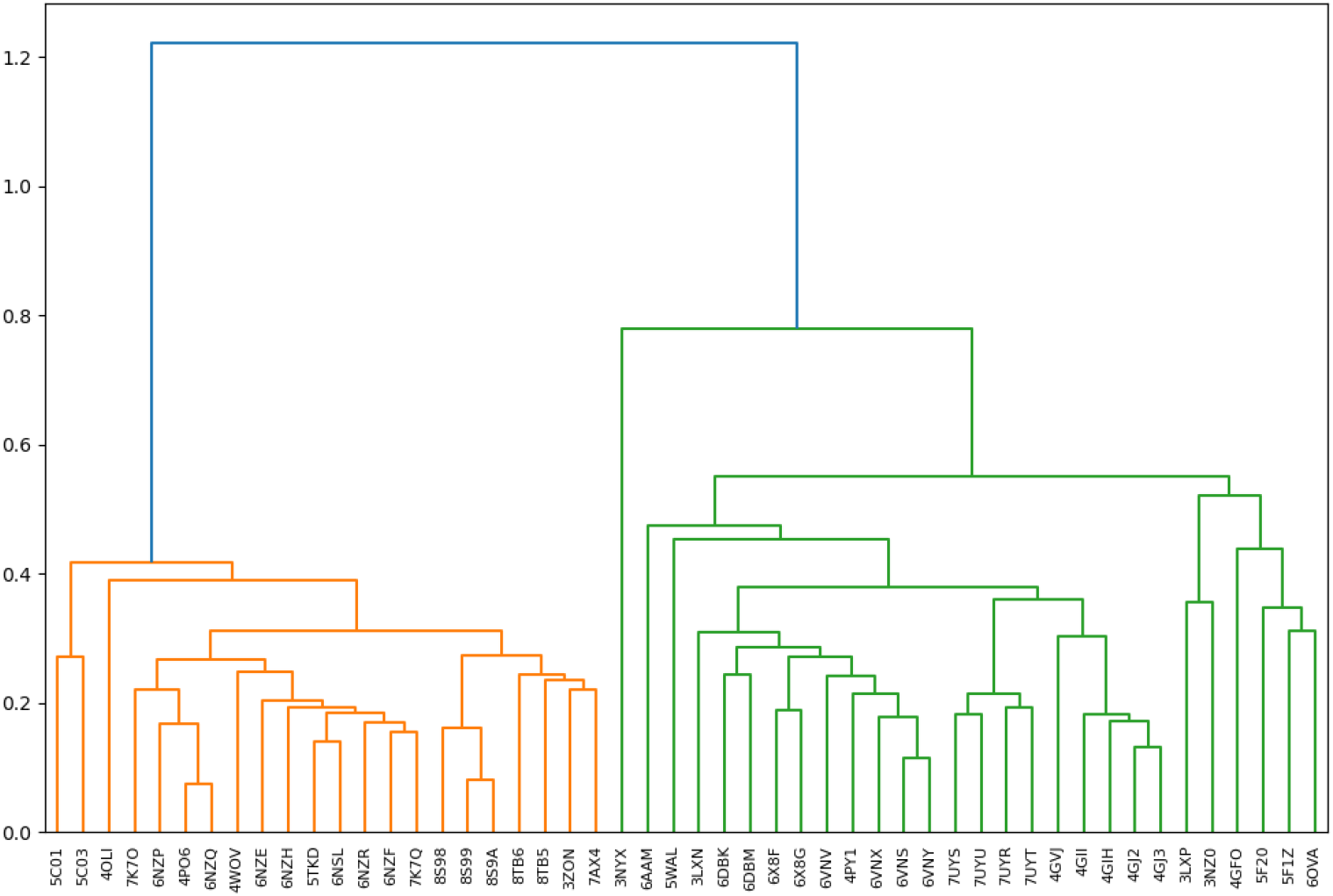
Dendrogram illustrating the results of hierarchical clustering.

**Figure 4:**
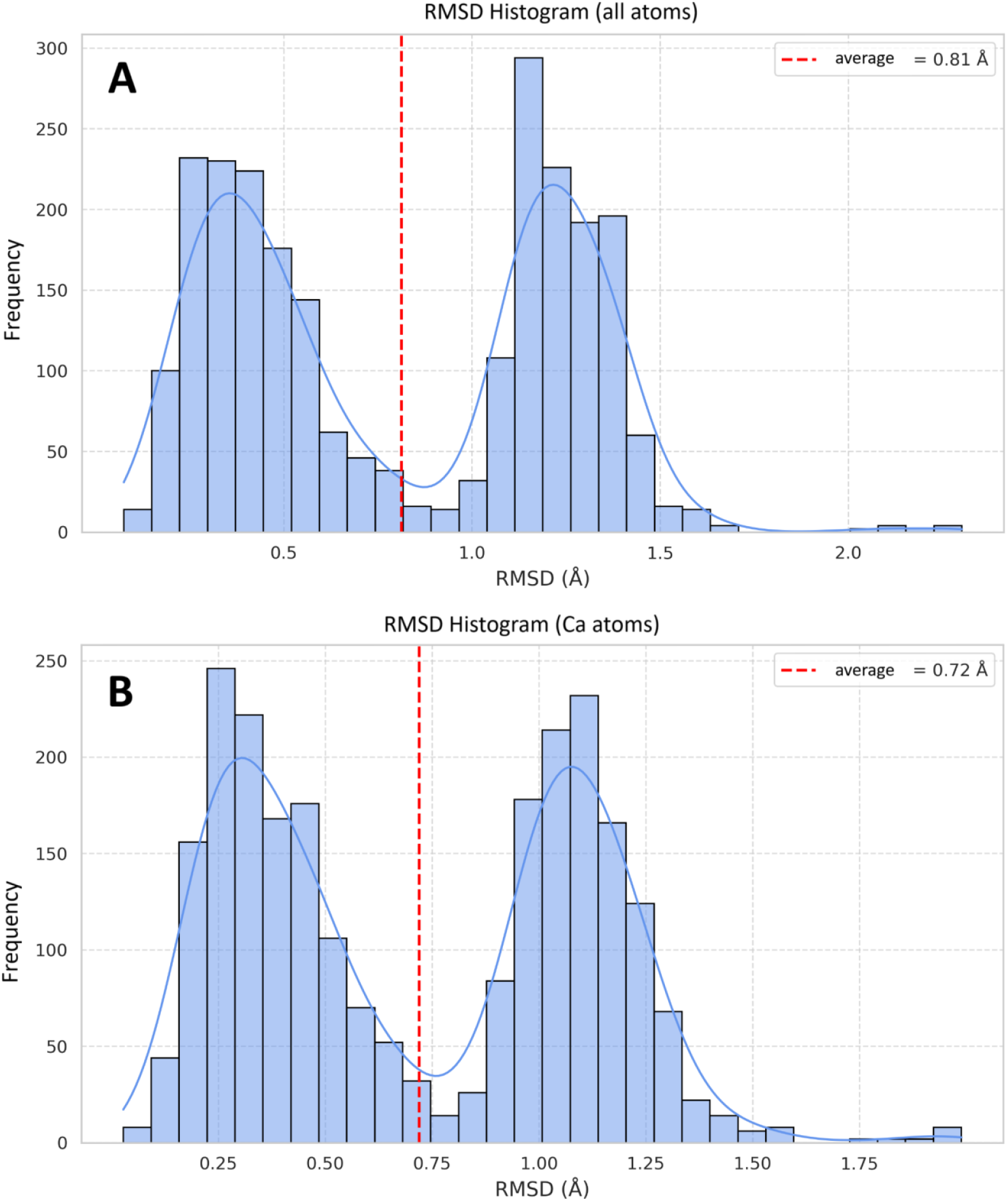
Histogram of RMSD distributions for all atoms (A) and Cα atoms (B).

Further analysis revealed that within each of these classes, there are subgroups of structures that are either closely related or noticeably different from one another. These differences are influenced both by the type of bound ligand and by specific structural variations in the protein. Some structures formed dense, homogeneous clusters, while others exhibited pronounced deviations, potentially indicating unique conformations or ligand-induced rearrangements. For example, structures with PDB IDs 8S98, 8S99, and 8S9A contain highly similar ligands sharing a common core scaffold (Figure 5), which explains their proximity and clustering within a single group.

**Figure 5:**
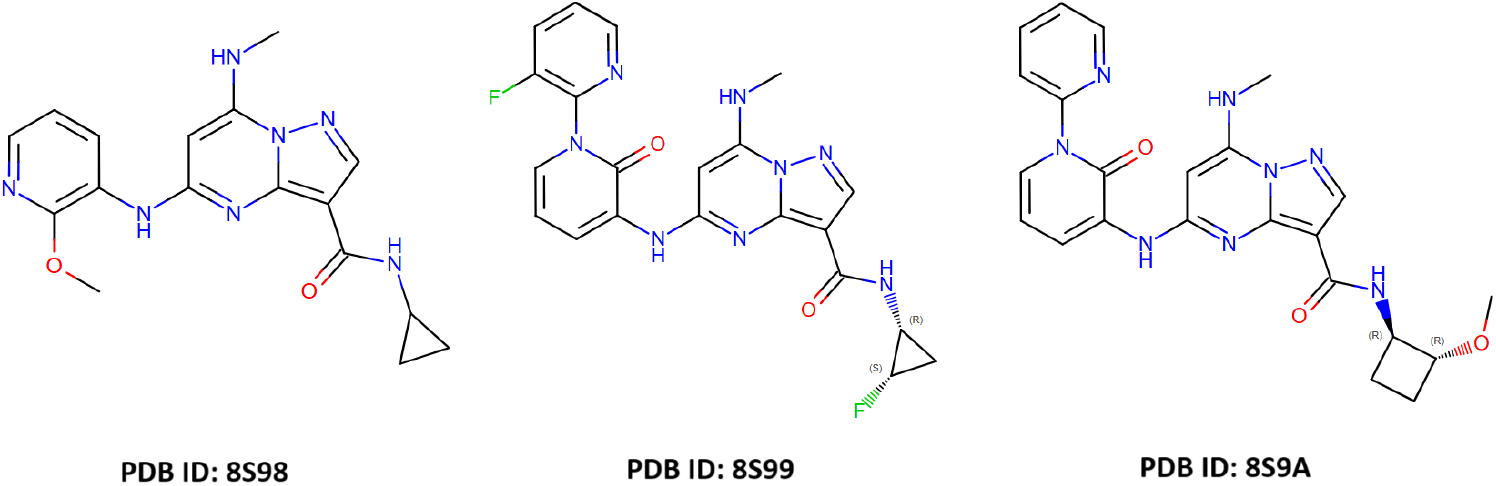
Comparison of ligands from PDB structures 8S98, 8S99, and 8S9A, highlighting their common scaffold and structural similarity.

## CONCLUSION

The developed tool, PocketMaster, demonstrated high efficiency in performing automated, scalable, and reproducible analyses of protein pocket structural similarity. Its flexible architecture - incorporating diverse pocket definition strategies, alignment methods, RMSD computation options, and clustering algorithms - enables adaptation to a broad spectrum of research tasks.

Application of PocketMaster to the analysis of TYK2 ligand-binding pocket variability revealed a distinct structural separation into two domains - kinase and pseudokinase - as well as conformational subtypes within these domains associated with ligand type and protein states. These findings not only confirm previously established structural characteristics of TYK2 but also underscore the potential of PocketMaster for data systematization and quantitative assessment of structural diversity, contributing to a deeper understanding of functionally relevant regions of the protein and supporting the selection of representative structures for drug development.

Thus, PocketMaster can serve as a powerful tool in the arsenal of structural bioinformaticians and drug designers, enabling deeper understanding of protein structural diversity, identification of stable conformations, and selection of promising target forms for rational drug design.

## DATA AND SOFTWARE AVAILABILITY

All data and software used in this study are freely available. The PocketMaster source code, together with example input files and documentation, can be accessed at https://github.com/narek-abelyan/PocketMaster

